# A structure-guided classification framework reveals the diversity and catalytic architecture of BECR ribonuclease

**DOI:** 10.64898/2026.08.28.747851

**Authors:** Kristi Pham, Gianlucca G. Nicastro, Audrey R. Long, L. Aravind, Claus O. Wilke, Robson Francisco de Souza, Ethel Bayer-Santos

**Affiliations:** Department of Molecular Biosciences, College of Natural Sciences, The University of Texas at Austin, Austin, USA; Division of Intramural Research, National Library of Medicine, National Institutes of Health, Bethesda, United States, USA; Department of Integrative Biology, College of Natural Sciences, The University of Texas at Austin, Austin, USA; Departamento de Microbiologia, Instituto de Ciências Biomédicas, Universidade de São Paulo, São Paulo, Brazil

**Keywords:** Biological conflict systems, Ribonuclease toxins, BECR fold, Protein evolution

## Abstract

Microorganisms across all domains of life engage in molecular conflict, deploying toxins to inhibit competitors or respond to biological threats. Among these, ribonuclease toxins are particularly widespread and diverse. A substantial fraction is associated with the BECR fold, a compact α/β architecture that supports RNase activity despite extensive divergence. Although several canonical members are well characterized, many BECR-fold proteins remain difficult to identify because of low sequence similarity, variation in catalytic residues, and structural elaborations that obscure evolutionary relationships. The growing availability of high-confidence protein structure predictions provides an opportunity to reassess this deeply divergent protein landscape. Here, we integrate iterative profile-HMM searches, profile-similarity networks, structural analyses, active-site mapping, and genomic context to examine BECR proteins across the tree of life. Our analysis resolves an expanded BECR-fold landscape comprising canonical BECR and BECR-like superfamilies, refines the organization of canonical BECR proteins and identifies previously unrecognized families. We further validate BECR-Tox2 as a toxin neutralized by a cognate immunity protein and show that its homologs occur in both Menshen-like anti-phage systems and polymorphic toxin loci. Together, these findings expand and clarify the BECR-fold landscape and provide a framework for identifying and interpreting highly divergent proteins of this fold.

## INTRODUCTION

Life at the microscopic scale is shaped by intense biological conflict, where organisms deploy diverse molecular strategies to secure resources, defend against biological threats, and compete with neighboring cells (Granato et al., 2019). These interactions are frequently mediated by toxins with broad functional diversity (Ruhe et al., 2020; Zhang et al., 2012), which are delivered through a variety of mechanisms and play central roles in shaping microbial community and ecological dynamics (Granato et al., 2019; Peterson et al., 2020).

A substantial fraction of these toxins function as nucleases and can be organized into protein superfamilies based on common ancestry, with conserved structural cores often providing an important signal of deep homology. Examples include the all-α helical HEPN (Higher Eukaryotes and Prokaryotes Nucleotide-binding) superfamily (Anantharaman et al., 2013) and the HNH (His-Me finger) superfamily (Jablonska et al., 2017). Among these, the BECR fold (named after the founding members Barnase, EndoU, Colicin E5/D, and RelE) represents a particularly broad structural framework that unifies diverse ribonucleases (Zhang et al., 2012). Representative members include Barnase, one of the earliest characterized ribonucleases (Meiering et al., 1992), RelE from toxin–antitoxin (TA) systems (Neubauer et al., 2009), colicins D and E5 (Ogawa et al., 2006; Tomita et al., 2000), and prokaryotic EndoU toxin (Michalska et al., 2018).

The BECR fold consists of a compact α/β RNase scaffold formed by an N-terminal α-helix followed by a four-stranded meandering β-sheet (Zhang et al., 2012). Comparative sequence and structural analyses first revealed that this shared architecture linked several previously unrelated families of RNA-targeting toxins, establishing the BECR fold as a coherent evolutionary framework (Zhang et al., 2012). Recognition of this conserved structural core subsequently enabled the discovery and functional interpretation of numerous toxin families despite little or no detectable sequence similarity (Cuthbert et al., 2022; Gucinski et al., 2019; Iyer et al., 2017; H. Li et al., 2023; Michalska et al., 2017, 2018a; Nicastro et al., 2026).

BECR proteins belong to the broader class of metal-independent ribonucleases. RNA cleavage proceeds through an internal transesterification reaction in which a general base activates the ribose 2′-OH, promoting nucleophilic attack on the adjacent phosphate and formation of a 2′,3′-cyclic phosphate (2′,3′-cP) intermediate, while a general acid protonates the departing 5′-oxygen (Shigematsu et al., 2018). Although this catalytic strategy is shared across characterized BECR ribonucleases, the identities of the catalytic residues vary substantially among families. Histidine frequently functions as a general acid or base, but glutamate, lysine, arginine, tyrosine, and other polar residues have also been implicated in catalysis or transition-state stabilization in different BECR proteins (Inoue-Ito et al., 2012; Lin et al., 2005; Neubauer et al., 2009; Takebe et al., 2024). This variability complicates the identification of catalytic residues and further obscures evolutionary relationships among highly divergent family members.

Despite these advances, the broader evolutionary organization of the BECR fold remains poorly understood. In addition to the low sequence similarity, many members of this group contain insertions, deletions, duplications, and permutations that obscure both sequence- and structure-based homology detection (Zhang et al., 2014). At the same time, BECR proteins preserve the same metal-independent ribonucleolytic chemistry while varying the identity of catalytic residues, a property observed in several highly divergent enzyme superfamilies (Hasson et al., 1998; Todd et al., 2001; Zhang et al., 2014). Together, these features complicate annotation, obscure evolutionary relationships, and hinder the development of a unified guideline capable of describing the full diversity of BECR proteins.

The growing availability of high-confidence protein structure predictions now provides an opportunity to revisit this problem. Large-scale structural resources have transformed the discovery and classification of deeply divergent protein families by enabling comparisons beyond the limits of sequence conservation. For the BECR fold, these resources make it possible not only to identify previously unrecognized proteins but also to reassess higher-order relationships across the superfamily and evaluate longstanding questions concerning its evolutionary diversification.

Here, we combine different approaches to reassess proteins associated with the BECR fold across the tree of life. By integrating this information, we define a unified classification framework for BECR-fold proteins. Our analyses reveal an expanded BECR landscape comprising canonical BECR and BECR-like superfamilies, refine the organization of canonical BECR proteins, support an evolutionary relationship between RNase A-like and EndoU-like proteins, and identify previously unrecognized toxin families. In addition, we provide an interactive resource of structures and sequence alignments to facilitate future identification and analysis of BECR proteins (https://kristiphammie.github.io/becr/). Together, these findings expand and clarify the BECR-fold landscape, improve the annotation of highly divergent RNase toxins, and provide a guideline for investigating their evolutionary relationships and biological functions.

## RESULTS

### Defining a representative dataset across the expanded BECR-fold landscape

To systematically capture the breadth of the BECR fold, we constructed a curated dataset using an explicit, criteria-based framework designed to recover both canonical and highly divergent members. Protein families were included if they satisfied at least one of three conditions: (i) prior annotation as a BECR-fold RNase; (ii) experimentally determined or predicted structural similarity to canonical BECR-fold proteins; or (iii) evidence of a shared ribonucleolytic mechanism generating a 2′,3′-cP intermediate. To ensure specificity, families included under the third criterion were additionally required to exhibit a recognizable BECR-fold core topology. This constraint enabled inclusion of highly divergent proteins while avoiding incorporation of unrelated RNases that share the same catalytic mechanism (Table S1).

Using these characteristics, we manually assembled a dataset by tracing the BECR literature. Starting with the original BECR study (Zhang et al., 2012), we followed citations to studies describing related toxins and iteratively expanded this reference network until no additional families could be identified. Representative alignments for each family were obtained from PFAM models (Mistry et al., 2021), original publications, or generated *de novo* when suitable alignments were unavailable. For PFAM- and publication-derived models, alignments that appeared to truncate conserved regions of the fold were manually revised and replaced with curated versions generated in this study. Both the original and curated alignments are available through our online resource (https://kristiphammie.github.io/becr/).

This process produced a dataset of 13,424 sequences representing 67 distinct protein families (Table S2A). These families span a broad range of biological contexts, including bacterial conflict systems, eukaryotic ribonucleases, and viral proteins (Table S1). Each family was represented by a single reference structure derived from an experimentally determined structure or a high-confidence AlphaFold prediction (Jumper et al., 2021) together with a corresponding multiple-sequence alignment for downstream analyses (https://kristiphammie.github.io/becr/). This curated dataset provides a representative view of currently known BECR-fold diversity and serves both as a reference for comparative analyses and as the seed set for the iterative sequence searches used to identify additional BECR families described below.

### Systematic expansion of families associated with the BECR fold

After establishing a representative dataset spanning the currently recognized diversity of the BECR fold, we sought to determine whether additional, previously unrecognized homologs could be identified through iterative sequence searches. Because BECR-fold proteins exhibit exceptionally low pairwise sequence similarity, conventional sequence-based approaches are likely to recover only a limited fraction of their diversity. We therefore implemented an iterative workflow that combined sensitive profile-HMM searches with sequence similarity network analysis (Fig. 1A). Curated multiple-sequence alignments representing each BECR-fold family were used to generate profile HMMs for JackHMMER searches (Eddy, 2011). Newly recovered homologs were then organized into sequence similarity networks (SSNs), allowing previously unsampled sequence clusters to be identified, curated into new alignments, and incorporated into subsequent rounds of profile construction and homology searches. These complementary approaches have been widely used to detect remote protein families and infer relationships among divergent proteins (Atkinson et al., 2009; Copp et al., 2019; Medini et al., 2006; Song et al., 2007).

**Fig 1.**
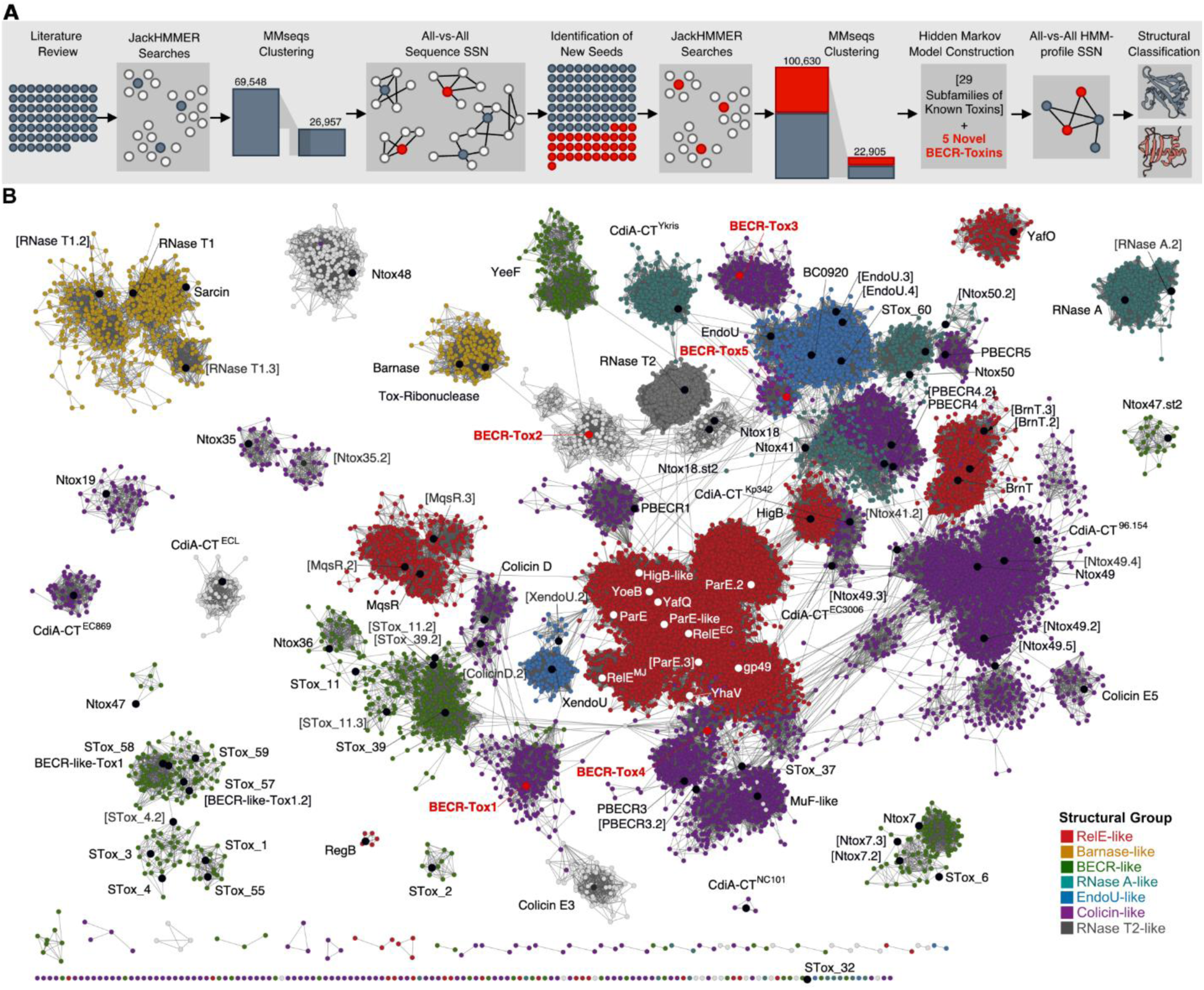
Sequence similarity network displays extensive sequence divergence and identifies new BECR families. (A) Pipeline schematic of SSN generation. (B) SSN of BECR fold proteins. Each node represents a protein sequence, and edges indicate pairwise similarity above an E-value threshold of 1e−2. Nodes are colored by structural grouping as shown in the legend. Black-labeled nodes denote seed sequences used as queries for homology searches for each family, while red labels indicate putative new families identified from the initial round of searches and included in the analysis.

In the first search, profile HMMs generated from the 67 curated BECR-fold families were used as queries for five rounds of JackHMMER searches against the UniRef50 database (Bateman et al., 2023; Eddy, 2011), using a relaxed inclusion threshold of E-value < 0.01 to recover distant homologs (Table S2B). These searches expanded the dataset from 13,424 to 69,547 sequences (Table S2C), which were reduced to 26,957 representative sequences after detection of redundant sequence groups (Table S2D). An SSN constructed from these representatives revealed a highly fragmented sequence landscape in which many BECR-fold families formed discrete clusters with little or no detectable connectivity to other families (Fig. S1). To systematically identify previously unrecognized diversity, we focused on clusters lacking representatives from the original 67 seed families. Representative sequences from 34 candidate clusters were manually curated into new multiple-sequence alignments and used to generate additional profile HMMs for a second round of JackHMMER searches using the same inclusion criteria (Table S2E). This second iteration expanded the dataset to 100,630 sequences (Table S2F), which were subsequently reduced to 22,905 representative sequences after redundancy reduction using less stringent parameters (Table S2G).

The final SSN organized this expanded dataset according to detectable pairwise sequence relationships and provided a sequence-level view of the BECR-fold landscape (Fig. 1B). Most established families formed discrete clusters, reflecting the extensive sequence divergence that characterizes the fold. At the same time, the SSN resolved previously unsampled clusters recovered through the iterative searches and enabled their relationship with the original seed families to be evaluated. Thus, the SSN served two purposes: it illustrated the fragmented organization of the BECR sequence space and provided the primary framework through which candidate new families were recognized.

We next evaluated whether the newly recovered clusters represented divergent subfamilies of established BECR-fold families or independent new families. Clusters retaining the domain architecture, sequence conservation patterns, predicted catalytic organization, and overall structural features of the family from which they were recovered were classified as subfamilies and named using the format *SeedName#*. In total, 31 new subfamilies of established families were identified. In contrast, clusters displaying distinct sequence signatures, structural features, or genomic contexts were classified as previously unrecognized BECR-fold families. Applying these criteria, we identified five families that were designated BECR-Tox1 through BECR-Tox5 (Fig. 1B). Among these, we focused on the experimental characterization of BECR-Tox2, which will be described below.

### Profile–profile network analysis resolves the higher-order organization of the BECR-fold landscape

Although the SSN resolved individual families and enabled the identification of previously unsampled sequence clusters, many families remained disconnected from the broader network because their pairwise sequence similarity fell below detectable levels (Fig. 1B). This fragmentation limited the ability of the SSN to resolve higher-order relationships across the expanded BECR-fold landscape. We therefore performed all-against-all comparisons of the curated family profiles and used the resulting similarities to construct a profile–profile network (Fig. 2). By integrating sequence information across entire protein families rather than comparing individual proteins, this analysis recovered remote similarities that were no longer detectable through pairwise sequence comparisons and reconnected many families that appeared isolated in the SSN. The two networks therefore provide complementary views of the expanded BECR landscape.

**Fig 2.**
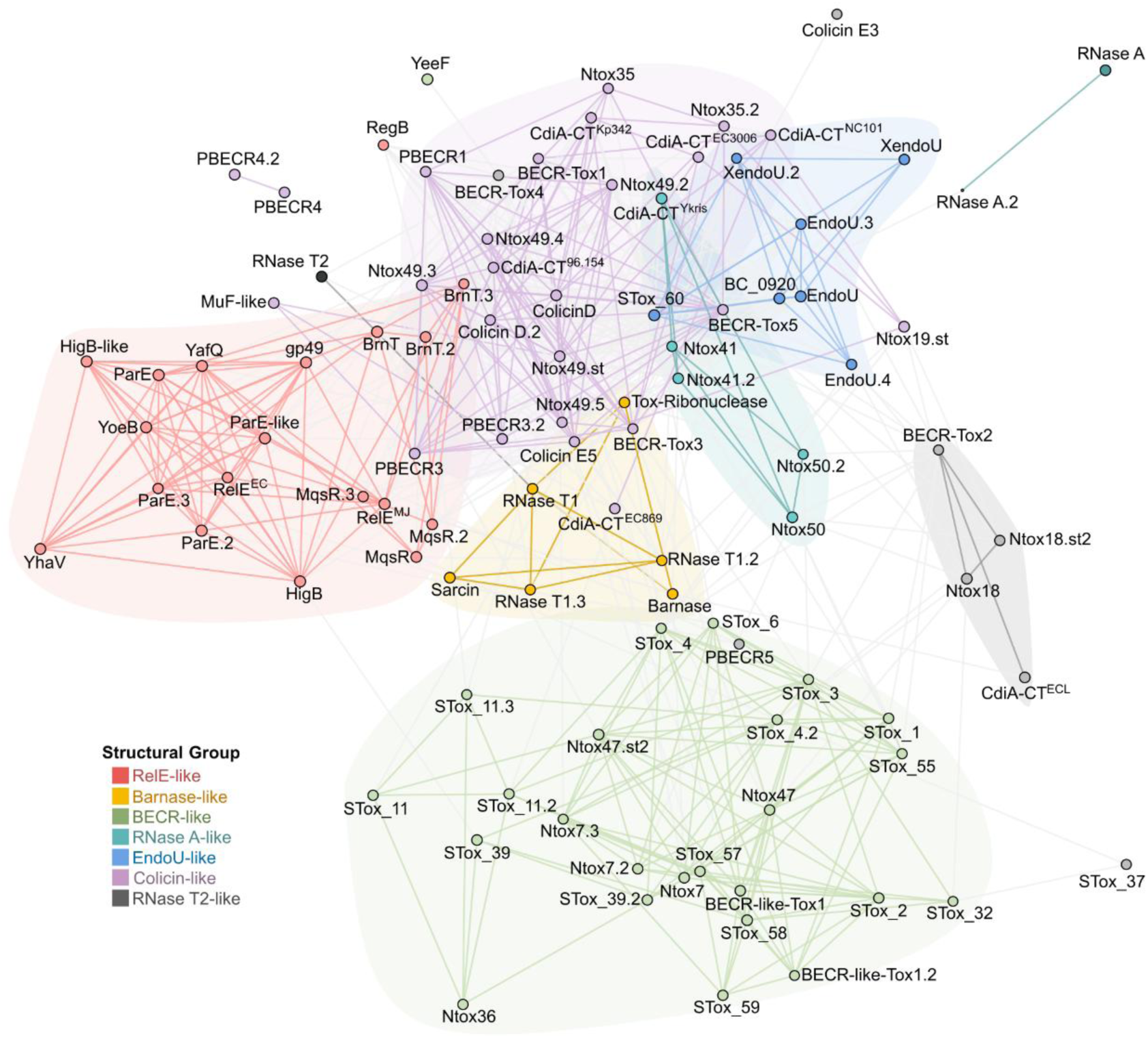
Profile–profile similarity network resolves two major BECR groups and reveals remote homology relationships. Each node represents an HMM profile, and edges denote significant profile– profile similarity based on HHpred comparisons with an E-value threshold of 1e-2. Node size is proportional to the number of sequences in the profile HMM and colored according to the structural classes defined in the subsequent section.

Inspection of the profile–profile network revealed an organization comprising two major groups (Fig. 2). The first encompasses the canonical BECR families initially described by Zhang et al. (2012) together with additional families subsequently assigned to this fold. The second, designated here as the BECR-like superfamily, includes several previously described toxin families (Nicastro et al., 2026; Zhang et al., 2012), and additional families recovered in the present analysis. We previously introduced the term BECR-like for a subset of these proteins based on shared structural features, but their evolutionary relationship remained unclear (Nicastro et al., 2026). The expanded profile–profile network reveals recurrent connections between canonical BECR and BECR-like families that were not apparent in the sequence-level SSN (Fig. 1B). Although these observations support a broader association between canonical BECR and BECR-like proteins, they do not conclusively distinguish deep homology from convergence.

To examine the internal organization of the network, we independently applied the Leiden and Louvain community-detection algorithms (Traag et al., 2019). Within the canonical BECR superfamily, the network resolves principal regions associated with the EndoU-like, RelE-like, and Colicin-like groups (Fig. 2). Smaller but coherent groups include the Barnase-like and BECR-Tox2-like groups. The placement of RNase A-like proteins within the EndoU-associated region further supports a relationship between these families that was difficult to recover from protein-level sequence comparisons alone. The network also clarifies the higher-order placement of several families whose assignments based on existing domain models or structural resemblance remained uncertain. The organization established here provided directions for the sequence, structural, and catalytic comparisons that follow.

### Structural and catalytic organization of the BECR and BECR-like superfamilies

We next examined whether the groups resolved by the profile–profile network are also distinguished by coherent structural and catalytic features. For each group, we compared the topology of representative structures, mapped conserved residues onto predicted active-site pockets, and evaluated the distribution of these features across family-specific alignments (Fig. 3 and Fig. 4). These analyses revealed recurrent structural configurations built around a compact BECR-fold core.

**Fig 3.**
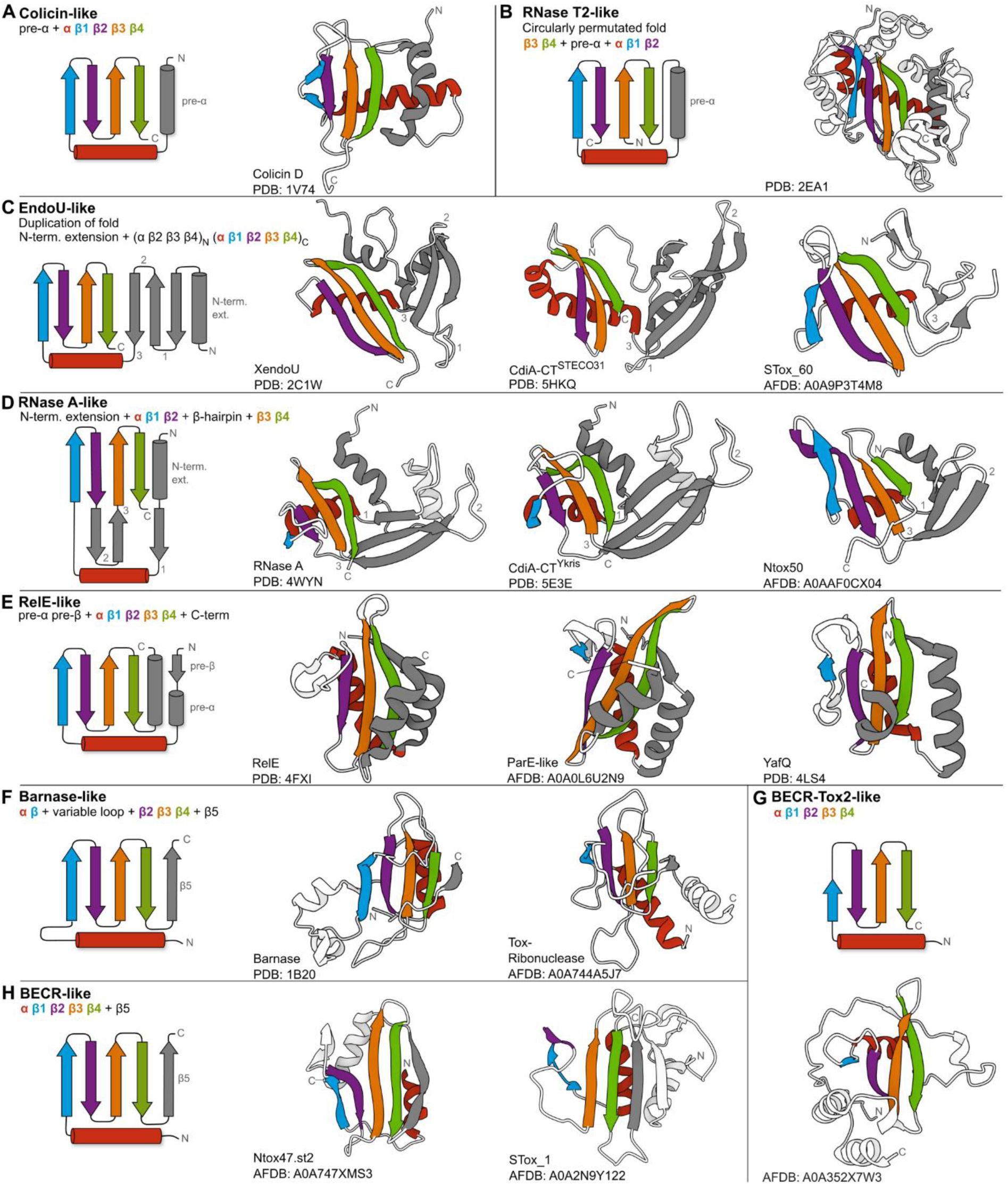
Structure-guided classification reveals distinct BECR-fold architectures built on a conserved core. Representative two-dimensional topologies (left) and corresponding three-dimensional structures (right) are shown for each BECR-fold group. Conserved core secondary-structure elements comprising the canonical BECR scaffold are colored as follows: α-helix (red), β1 (blue), β2 (purple), β3 (orange), β4 (green), whereas class-specific insertions, extensions, and auxiliary structural elements are shown in grey. Labels beneath each class indicate the structural nomenclature used throughout the manuscript. (A) Colicin-like group. (B) RNase T2-like represents a circularly-permutated Colicin-like. (C) EndoU-like group. (D) RNase A-like group. Numbering in panels (C) and (D) indicates differences in linker connectivity between conserved core secondary-structure elements. (E) RelE-like group. (F) Barnase-like group. (G) BECR-Tox2-like group. (H) BECR-like class.

**Fig 4.**
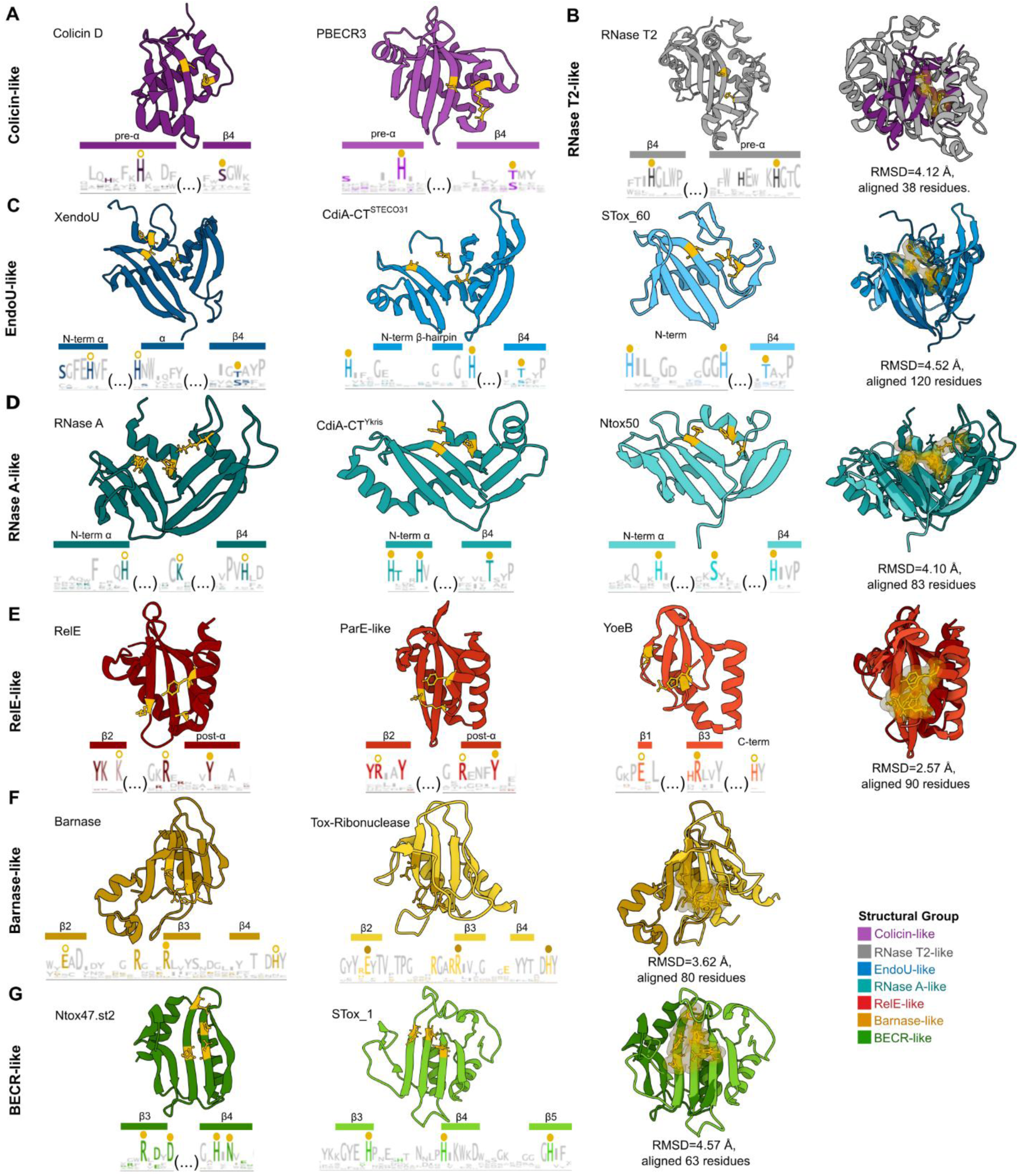
BECR groups share a conserved active site position despite variation in catalytic residues. Representative structures are shown with conserved residues highlighted in yellow (ball-and-stick), above sequence logos indicating residue conservation; yellow markers denote residues corresponding to those mapped onto the structures. Structural superpositions (TM-align) of representative members are shown on the right, illustrating the conserved spatial positioning of the active site across subclasses in a molecular surface representation.

#### The canonical BECR superfamily

Within the canonical BECR superfamily, the profile–profile network resolves five principal groups corresponding to the Colicin-like, EndoU-like, RelE-like, Barnase-like, and BECR-Tox2-like groups (Fig. 2). Each comprises multiple protein families connected by profile-profile similarity and supported by characteristic structural or active-site features. The structural comparisons below also consider two special configurations that warrant separate treatment. RNase A-like proteins are associated with the EndoU-like group but exhibit distinct strand connectivity, whereas RNase T2 adopts a circularly permuted version of the topology found in Colicin-like proteins. Together, these comparisons define the principal structural and predicted catalytic configurations of the canonical BECRs and provide a reference for evaluating the more divergent BECR-like proteins.

##### Colicin-like

The Colicin-like group is distinguished by a pre-α helix positioned immediately before the canonical BECR core and arranged parallel to the terminal β4-strand, yielding the characteristic topology [pre-α + αβ1β2β3β4] (Fig. 3A). Representatives span a continuum from architectures that closely resemble the inferred canonical BECR scaffold to highly modified forms that retain only a subset of the core secondary-structure elements. Canonical members such as colicins D and E5 preserve the complete topology, whereas several phage-associated BECR proteins (e.g. PBECRs) and contact-dependent inhibition (CDI) toxins display progressive remodeling of the scaffold through reduction or loss of β1 and incorporation of additional β-strands preceding the pre-α element (https://kristiphammie.github.io/becr/). This trend culminates in the CDI toxin CdiA-CT^EC869^, which retains only β3 and β4 from the canonical four-stranded sheet while preserving an overall BECR architecture (Wang et al., 2022).

Despite this structural variability, the predicted active-site architecture is comparatively conserved. Conserved active-site residues are typically positioned along the pre-α element and terminal β4, forming a pocket that depends strongly on the group-specific pre-α extension (Fig. 3A and Fig. 4A). A conserved histidine frequently occupies a position consistent with a catalytic role in acid-base chemistry, as observed in experimentally characterized representatives such as colicin D and CdiA-CT^96.154^ (Graille et al., 2004; Jones et al., 2017). Additional conserved lysine or arginine residues commonly occupy nearby positions within the pocket and may contribute to catalysis, substrate positioning, or transition-state stabilization (Table S1). Notably, the recurrent reduction or loss of β1 in many Colicin-like proteins occurs outside the active-site region, suggesting that this element is dispensable for maintaining catalytic geometry.

Functionally characterized members of the Colicin-like group predominantly target tRNAs, often with isoacceptor-level specificity (Gucinski et al., 2019; Jones et al., 2017; Ogawa et al., 2006, 2021; Wang et al., 2022). These proteins are widely deployed in biological conflict systems. The broad structural variability observed within the group, combined with conservation of the active-site region, suggests that many of the lineage-specific structural additions surrounding the BECR core may contribute to substrate recognition or delivery rather than to the catalytic reaction itself.

The structural framework established by the Colicin-like group also clarifies several ambiguous domain classifications. DUF4258 was recently assigned to the RelE/ParE superfamily (Gerdes, 2025), yet its domain boundaries substantially overlap those of the previously described Ntox49 family (Zhang et al., 2012). Consistent with this observation, representatives assigned to DUF4258 consistently adopt the characteristic Colicin-like topology and cluster with Colicin-like proteins in both sequence- and profile-based analyses (Fig. 1 and Fig. 2). This data therefore supports placement of DUF4258 within the Colicin-like branch of the canonical BECR superfamily.

A similar reclassification emerged for the poorly characterized PFAM family DUF6883. Our new BECR-Tox3 profile frequently recovers proteins annotated as DUF6883, indicating substantial overlap between the two models. Gene-neighborhood analysis further revealed that BECR-Tox3/DUF6883 genes are commonly associated with a conserved downstream DUF4926 protein, suggesting a toxin-immunity organization. In addition, BECR-Tox3/DUF6883 domains frequently occur as C-terminal fusions to proteins containing an N-terminal Phage_Mu_F domain. Previous studies identified related MuF-associated PBECR proteins as phage-encoded BECR effectors predicted to function as RNases (Iyer et al., 2017). The recurrent association of DUF6883 with both DUF4926 and Phage_Mu_F architectures therefore supports its assignment to the BECR-Tox3 family and places this group within a previously recognized repertoire of phage-associated BECR effectors.

##### RNase T2-like

RNase T2 was proposed to represent a circularly permuted member of the BECR superfamily (H. Li et al., 2023). Unlike the network-defined groups described above, RNase T2 warrants separate consideration because its relationship to canonical BECR proteins is obscured by extensive rearrangement of the primary sequence. Despite minimal detectable sequence similarity and substantial topological reorganization, the overall structure of RNase T2 preserves the defining architectural features of the BECR fold (Fig. 3B).

Li *et al*. (2023) showed that manual reconstruction of the permuted sequence reveals similarities that are otherwise masked by the altered connectivity of the fold. Consistent with this proposal, our structural comparisons identify a particularly strong correspondence between RNase T2 and members of the Colicin-like group (Fig. 2). Most notably, the conserved histidine located within the pre-α element of RNase T2 occupies the same structural position as the active-site histidine found in Colicin-like proteins (Fig. 3B and Fig. 4B). This residue is preceded by a semi-conserved lysine that is also observed in several Colicin-like representatives, including colicin D, PBECR3, and CdiA-CT^EC869^. A second conserved histidine is positioned on the terminal β4, mirroring the organization observed in Colicin-like proteins.

The preservation of both active-site architecture and catalytic residue positioning is notable given the extensive topological rearrangement that separates RNase T2 from canonical BECR proteins. Although circular permutation has altered the connectivity of the fold, the catalytic framework remains largely intact. These observations support the interpretation that RNase T2 represents a highly diverged derivative of the Colicin-like group rather than an independently evolved lineage.

##### EndoU-like

The EndoU-like group is distinguished by a duplication in the BECR core, yielding the topology [(αβ2β3β4)_N_ + (αβ1β2β3β4)_C_] (Fig. 3C). Consistent with the model originally proposed by Zhang et al. (2012), this arrangement can be interpreted as an N-terminal duplication of the canonical BECR core, although the duplicated unit frequently exhibits varying degrees of reduction. EndoU-like proteins from bacterial, eukaryotic, and viral systems display substantial architectural variation. Representatives including XendoU, BC_0920, and CdiA-CT^STECO31^ retain the duplicated framework but display marked variability in the N-terminal module, ranging from near-canonical BECR cores in CdiA-CT^STECO31^ to strongly reduced modules in STox_60 (Fig. 3C). By contrast, the C-terminal BECR module is conserved across the group, indicating that it forms the principal structural anchor while the duplicated N-terminal module is more variable.

This structural asymmetry is reflected in the organization of the active site. EndoU-like proteins exhibit one of the most conserved active-site arrangements within the canonical BECR superfamily, typically retaining two histidine residues together with a semiconserved polar residue, usually serine or threonine, positioned on the terminal β4 (Fig. 3C and Fig. 4C). Thus, the predicted active site is assembled from residues contributed by both the N-terminal variable module and the conserved C-terminal core. This arrangement persists even in structurally reduced members such as STox_60, indicating that active-site organization remains constrained despite substantial variation in the modules.

Functionally characterized EndoU-like proteins act on diverse RNA substrates. XendoU and viral NendoU proteins cleave single- or double-stranded RNAs, frequently with a preference for uridine-containing sequences (Gioia et al., 2005; Kim et al., 2020; Renzi et al., 2006) whereas bacterial EndoU-like toxins target tRNA and rRNA (Michalska et al., 2018). Thus, substantial differences in substrate preference and biological function occur within this group. The broad distribution of EndoU-like proteins across cellular and viral systems further indicates that this architecture has been recruited into diverse biological contexts.

##### RNase A-like

RNase A-like proteins are distinguished by a β-hairpin inserted between β2 and β3 of the BECR core. In several members, this hairpin pairs with a β-strand preceding the core α-helix to form an auxiliary β-sheet (Fig. 3D). Although this arrangement has a different strand connectivity from the duplicated architecture of EndoU-like proteins, the auxiliary β-sheet occupies a comparable spatial position. RNase A-like proteins have not previously been included within the BECR fold because of their divergent topology and the absence of detectable sequence similarity to canonical BECRs (Cuthbert et al., 2022). In our profile–profile network, however, multiple RNase A-like families consistently fall within the EndoU-associated region (Fig. 2). This assignment is supported by structural comparisons showing similar spatial organization of the core and auxiliary elements in these two groups (Fig. 3C and 3D).

RNase A-like proteins also share the general active-site organization observed in EndoU-like proteins, although the identities and structural origins of the contributing residues differ (Fig. 3C-D and Fig. 4C-D) (Batot et al., 2017; Jamet et al., 2015). An N-terminal histidine is frequently retained, consistent with previous evidence for an evolutionary relationship between the RNase A and EndoU families (Mushegian et al., 2020). By contrast, the serine or threonine associated with the terminal β4 in EndoU-like proteins is replaced by a second histidine in characterized RNase A-like proteins, while an additional catalytic residue is contributed by a variable loop (Cuchillo et al., 2011). In both groups, the active site is therefore assembled from a combination of core and auxiliary structural modules.

##### RelE-like

The RelE-like group is distinguished by an N-terminal insertion composed of a shortened β-strand and α-helix aligned parallel to the terminal β4, along with a C-terminal extension containing conserved active-site residues, yielding the topology [pre-β + pre-α + αβ1β2β3β4 + C-term] (Fig. 3E). RelE-like proteins exhibit the most conserved overall architecture among the canonical BECRs, with comparatively limited variation surrounding the core. The C-terminal region contains a flexible α-helix or loop that has been implicated in substrate recognition and catalytic function (Francuski & Saenger, 2009; G.-Y. Li et al., 2009). As observed in the Colicin-like group, several representatives lack β1 (e.g. RelE^EC^, RegB, and YhaV), indicating that this strand is not essential for maintaining the overall active-site architecture.

RelE-like toxins are predominantly ribosome-dependent mRNA interferases that cleave mRNA positioned in the ribosomal A site, often with codon-specific preferences mediated by interactions with the substrate and surrounding ribosomal RNA (Schureck et al., 2015; Harms et al., 2018; Gerdes, 2025). Consistent with this mode of action, conserved active-site residues are concentrated around β3, with additional contributions from β2 and the group-specific C-terminal region (Fig. 3E and Fig. 4E). A conserved C-terminal tyrosine frequently occupies the center of the active-site pocket and has been implicated in substrate stacking interactions and noncanonical catalytic mechanisms involving surrounding basic residues (Neubauer et al., 2009).

Unlike Colicin-like proteins, which typically rely on a single conserved positive residue to stabilize the transition state, RelE-like toxins display a broad distribution of positively charged residues across their surface. This expanded electrostatic environment is consistent with the extensive interactions required for ribosome binding and engagement of the mRNA substrate within the A site (Francuski & Saenger, 2009). Many RelE-like proteins lack a conserved histidine but nonetheless maintain a similar active-site geometry through alternative arrangements of basic and polar residues occupying equivalent structural positions (Dunican et al., 2015; Griffin et al., 2013). In members that do retain a histidine (e.g., YoeB- and HigB-like toxins) (Kamada & Hanaoka, 2005; Schureck et al., 2016), the residue occupies the same region of the active-site pocket, further supporting the idea that conservation of active-site organization is maintained despite variation in residue identity (Fig. 4E).

In contrast to several other canonical BECR groups, which are frequently associated with secreted toxins involved in intercellular antagonism, RelE-like proteins are predominantly associated with intracellular TA systems and related stress-response pathways (LeRoux & Laub, 2022). The RelE-like group therefore illustrates how the BECR scaffold can be adapted for a distinct biological role.

##### Barnase-like

The profile-profile network resolved a small but distinct group comprising Barnase, the fungal ribonucleases RNase T1 and α-sarcin (Lacadena et al., 1999; Loverix & Steyaert, 2001). Members of this group are characterized by a five-stranded β-sheet architecture in which a prominent C-terminal β5 extends the canonical four-stranded BECR core (Fig. 3F). Most also contain a variable insertion between β1 and β2, which forms an extended loop in some proteins, particularly α-sarcin. Although the original description of the BECR superfamily noted the presence of a short and variable fifth strand in some members (Zhang et al., 2012), our expanded analysis identifies numerous proteins with a well-developed β5 strand, indicating that expansion of the central β-sheet is a recurrent feature of the Barnase-like group rather than an occasional structural elaboration.

The active-site architecture of the Barnase-like group is likewise distinctive (Fig. 4F). Barnase provides the best-characterized example, in which a glutamate located near the middle of β2 functions as the general base and a histidine positioned between β4 and β5 functions as the general acid (Meiering et al., 1992, 1993). Consistent with this arrangement, a glutamate occupying an equivalent structural position is conserved across many members of the group (Fig. 4F). The variable β1-β2 insertion lies immediately adjacent to the catalytic region and may contribute to substrate recognition or specificity.

##### BECR-Tox2-like

The profile–profile network resolved our newly identified family, BECR-Tox2, together with other toxins as a distinct group within the canonical BECR superfamily (Fig. 2). Although members of this group retain the canonical BECR topology, they are distinguished by a conserved N-terminal extension preceding the core α-helix that is absent from other canonical BECR families (Fig. 3G). Despite exhibiting little predicted secondary structure, this N-terminal region consistently contributes a conserved KH motif to the active-site region. The histidine residue of this motif is positioned adjacent to a second conserved histidine located near the N-terminus of β4, forming the characteristic catalytic arrangement shared across the group. These features define a distinctive structural and catalytic signature that unifies the BECR-Tox2-like lineage. Because BECR-Tox2 is representative of this lineage and occurs in multiple biological conflict contexts, we selected it for experimental characterization, which is described further below.

#### The BECR-like superfamily

The profile-profile network resolved a second major group associated with the expanded BECR fold, designated here as the BECR-like superfamily (Fig. 2). We previously introduced the term BECR-like for a subset of these proteins based on structural similarities to canonical BECR proteins (Nicastro et al., 2026), but their broader organization and relationship with the canonical BECR superfamily remained unresolved.

For structural comparisons, we designated secondary-structural elements according to their topological correspondence with the canonical BECR core. Despite substantial variation across families, the most consistently conserved arrangement comprises a central α-helix followed by β3- and β4-equivalent strands separated by a characteristically extended loop and a C-terminal β5 that is broadly conserved throughout the superfamily (Fig. 3H). In several members, an additional N-terminal β-strand precedes the α-helix and packs against β5. By contrast, elements corresponding to canonical β1 and β2 vary considerably and are frequently reduced or absent. This structural organization is mirrored in the distribution of conserved active-site residues, which are typically positioned at the C-terminal end of the β3-equivalent strand and the N-terminal end of the β4-equivalent strand, flanking the extended β3-β4 loop (Fig. 4G). Thus, despite extensive variation in the β1- and β2-equivalent regions, the α-helix-β3-equivalent-loop-β4-equivalent-β5 arrangement defines a common structural and catalytic organization across the BECR-like superfamily.

The extreme divergence of the BECR-like superfamily makes its evolutionary relationship to canonical BECR proteins difficult to establish with confidence. Conventional structure-similarity searches initiated with BECR-like representatives fail to recover canonical BECR proteins, and pairwise sequence comparisons reveal no detectable similarity between the two groups. Although profile-profile analyses recover connections between BECR-like and canonical BECR families, these matches are generally weak. However, the similarities between BECR-like and canonical BECR proteins are recurrent. Multiple BECR-like families recover profile-level matches to several canonical BECR groups, forming a coherent network-level pattern rather than a single connection. In addition, BECR-like proteins frequently occur in multidomain, and genomic contexts occupied by canonical BECRs in related biological conflict systems and display a comparable organization of the predicted catalytic pocket, including recurrent use of histidine residues in positions consistent with catalytic roles. Taken together, these observations are more consistent with deep divergence from a common ancestral scaffold than with multiple independent origins, although the available evidence remains insufficient to distinguish conclusively between homology and convergence. We therefore use the designation BECR-like to recognize this coherent superfamily without treating its evolutionary relationship to canonical BECR proteins as resolved.

The assignment of the Ntox47 family within the BECR-like superfamily also clarifies a longstanding annotation issue. A representative member of this family is currently assigned to the PFAM family PF15657 and annotated as an HNH nuclease belonging to the broader EndoVII superfamily (Aravind et al., 2000; Zhang et al., 2011). However, our structural analyses indicate that PF15657 neither adopts an HNH fold nor resembles the canonical EndoVII architecture. Instead, it displays the defining structural and catalytic features of the Ntox47-like branch of the BECR-like superfamily. These observations indicate that PF15657 is misannotated and should be reassigned to the BECR-like superfamily. We therefore propose renaming this toxin to BECR-like-Tox1 (Fig. 2).

### BECR-Tox2 is associated with antiviral defense and polymorphic toxins systems

Among the newly identified families, BECR-Tox2 was selected for experimental characterization because it represents the founding member of a previously unrecognized group of canonical BECR proteins resolved by our profile-profile network analysis (Fig. 2). Members of this BECR-Tox2-like group retain the canonical BECR topology but are distinguished by a conserved N-terminal extension containing a characteristic KH motif that contributes to the predicted active-site region together with a second conserved histidine near β4 (Fig. 3G and Fig. 5A). These features define a distinct structural and catalytic configuration not observed in other canonical BECR lineages. Thus, characterization of BECR-Tox2 not only validates a new toxin family but also provides experimental insight into an entire branch of the canonical BECR superfamily.

**Fig 5.**
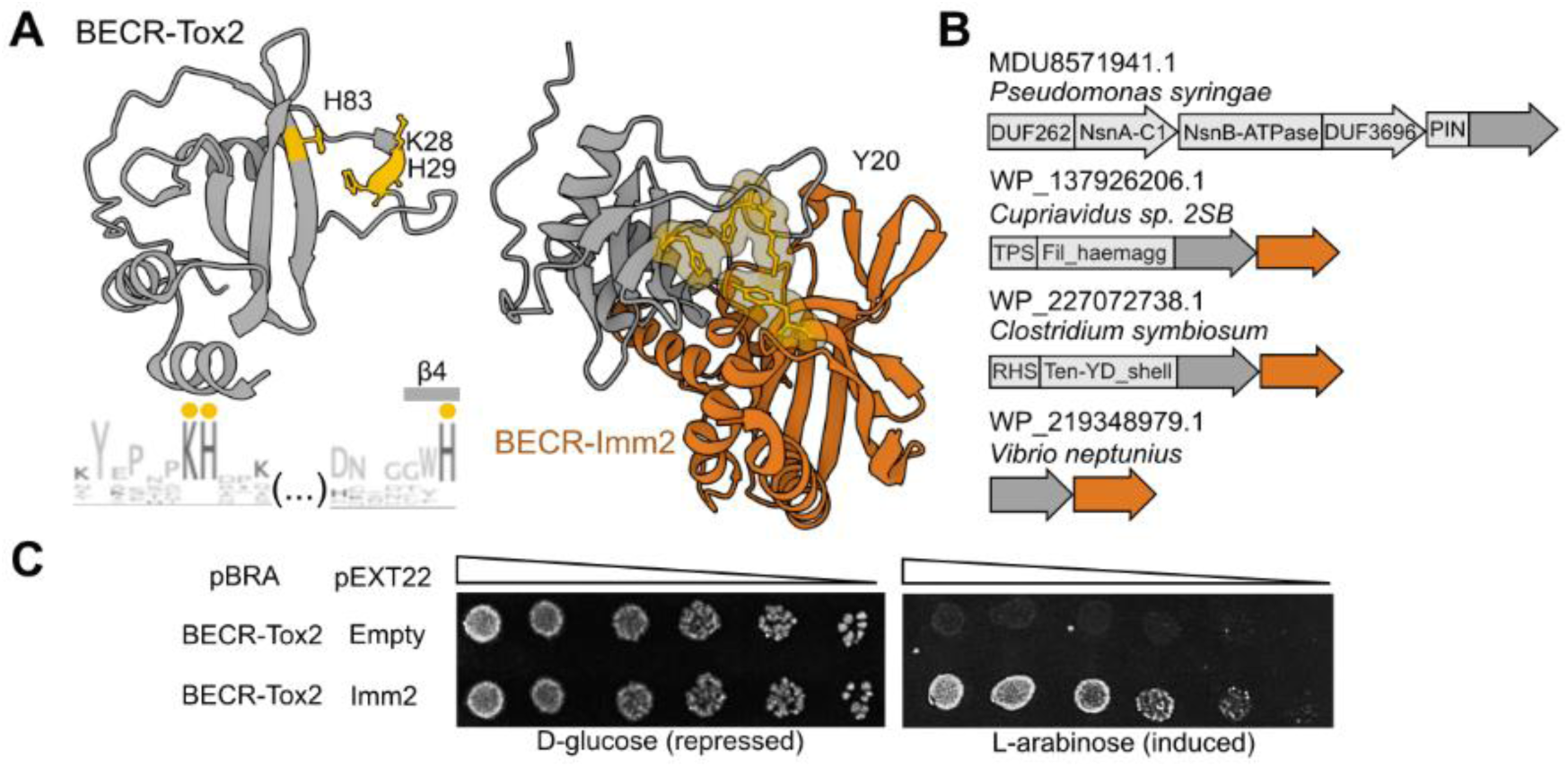
BECR-Tox2 functions as a toxin in antiviral defense and polymorphic toxin systems. (A) Left: Topology diagram of BECR-Tox2 with elements color-coded according to the conserved BECR core. The corresponding three-dimensional structure is shown with catalytic residues highlighted in ball-and-stick and indicated above the family logo (yellow markers). Right: AlphaFold-Multimer model of the BECR-Tox2–BECR-Imm2 complex, showing the predicted interaction mediated by Y20 of BECR-Imm2 (orange). (B) Representative gene neighborhoods illustrating the diversity of genomic contexts in which BECR-Tox2 occurs. Dark grey arrows denote BECR-Tox2 homologs, orange indicates the cognate immunity gene, and light grey represents surrounding genes. (C) Toxicity assay demonstrating the activity of BECR-Tox2.

Genomic neighborhood analysis revealed that most BECR-Tox2 homologs occur within the recently described Menshen antiphage defense system (Fig. 5B). In these loci, BECR-Tox2 is present as the C-terminal domain of a multidomain protein containing an N-terminal PIN nuclease domain and is associated with a ParB-family protein and an ABC ATPase (Krishnan et al., 2020; H. Li et al., 2025). Previous studies established that Menshen systems recruit diverse nuclease effectors, including members of the broader BECR superfamily, as interchangeable toxic outputs. However, BECR-Tox2 family members identified here were not described in those studies. Instead, our analyses indicate that Menshen systems contain multiple evolutionarily distinct BECR effectors and identify BECR-Tox2 as a previously unrecognized member of this repertoire. Comparative analyses further show that the conserved ParB-ABC ATPase module can associate with alternative nuclease domains, including BECR-, Toprim-, and HET-C-family proteins, suggesting that these domains function as interchangeable effector modules within a common defense architecture.

Although previous studies proposed that the N-terminal PIN domain associated with BECR domain is catalytically inactive due to the absence of canonical PIN catalytic residues (Gerdes, 2025; H. Li et al., 2025), our analyses do not support this interpretation. The PIN domain retains a conserved constellation of acidic and polar residues characteristic of active PIN nucleases, including residues occupying positions consistent with a canonical metal-binding active site. These observations raise the possibility that Menshen-associated BECR-Tox2 proteins function as dual nuclease effectors containing both PIN and BECR catalytic domains, although the biochemical roles of each domain remain to be determined.

While most BECR-Tox2 homologs occur in Menshen-like defense systems, a distinct subset is associated with genomic architectures characteristic of polymorphic toxin systems (Fig. 5B). In these loci, BECR-Tox2 is encoded adjacent to a small downstream gene lacking recognizable conserved domains. Based on this conserved association, we designated this protein BECR-Imm2 and hypothesized that it functions as a cognate immunity protein. Structural modeling using AlphaFold-Multimer predicted formation of a stable complex between a representative toxin-immunity pair from *Vibrio neptunius*, with BECR-Imm2 positioned to occlude the predicted active-site region of the toxin (Fig. 5A; Fig. S2). This arrangement is consistent with toxin neutralization through direct active-site inhibition, a common mechanism in polymorphic toxin systems (Alexei et al., 2024; Bartoli et al., 2023; Bosch et al., 2023; Liu et al., 2026).

To determine whether BECR-Tox2 functions as a toxin, we synthesized and expressed the *V. neptunius* homolog in *E. coli*. Expression of BECR-Tox2^VN^ produced a strong growth defect, demonstrating that the protein is toxic when expressed intracellularly (Fig. 5C). Co-expression of the associated BECR-Imm2^VN^ protein completely restored growth, confirming that BECR-Imm2 functions as a specific inhibitor of BECR-Tox2-mediated toxicity.

Together, these results establish BECR-Tox2 as the founding member of a previously unrecognized canonical BECR lineage associated with two major classes of biological conflict systems. Most homologs occur as candidate effectors within Menshen antiviral defense modules, whereas a distinct subset has been incorporated into polymorphic toxin systems together with dedicated immunity proteins. The occurrence of closely related BECR-Tox2 proteins in both contexts demonstrates that this newly defined BECR lineage has been repeatedly recruited into distinct conflict strategies, highlighting the evolutionary mobility and functional versatility of the BECR fold.

### BECR RNases are distributed across all domains of life

To assess the taxonomic distribution of BECR-fold proteins, we mapped all identified families across the tree of life using a nonredundant set of representative sequences (see Methods). This analysis provides an independent view of BECR diversity by examining the phylogenetic distribution of each family across bacterial, archaeal, eukaryotic, and viral lineages. BECR proteins are broadly distributed, with representatives detected in all domains of life (Fig. 6). However, this distribution is highly uneven across structural classes and families. RelE-like and Colicin-like proteins are widespread across multiple bacterial and archaeal phyla, whereas other classes are restricted to narrower taxonomic groups, reflecting lineage-specific expansions and losses (Fig. 6).

**Fig 6.**
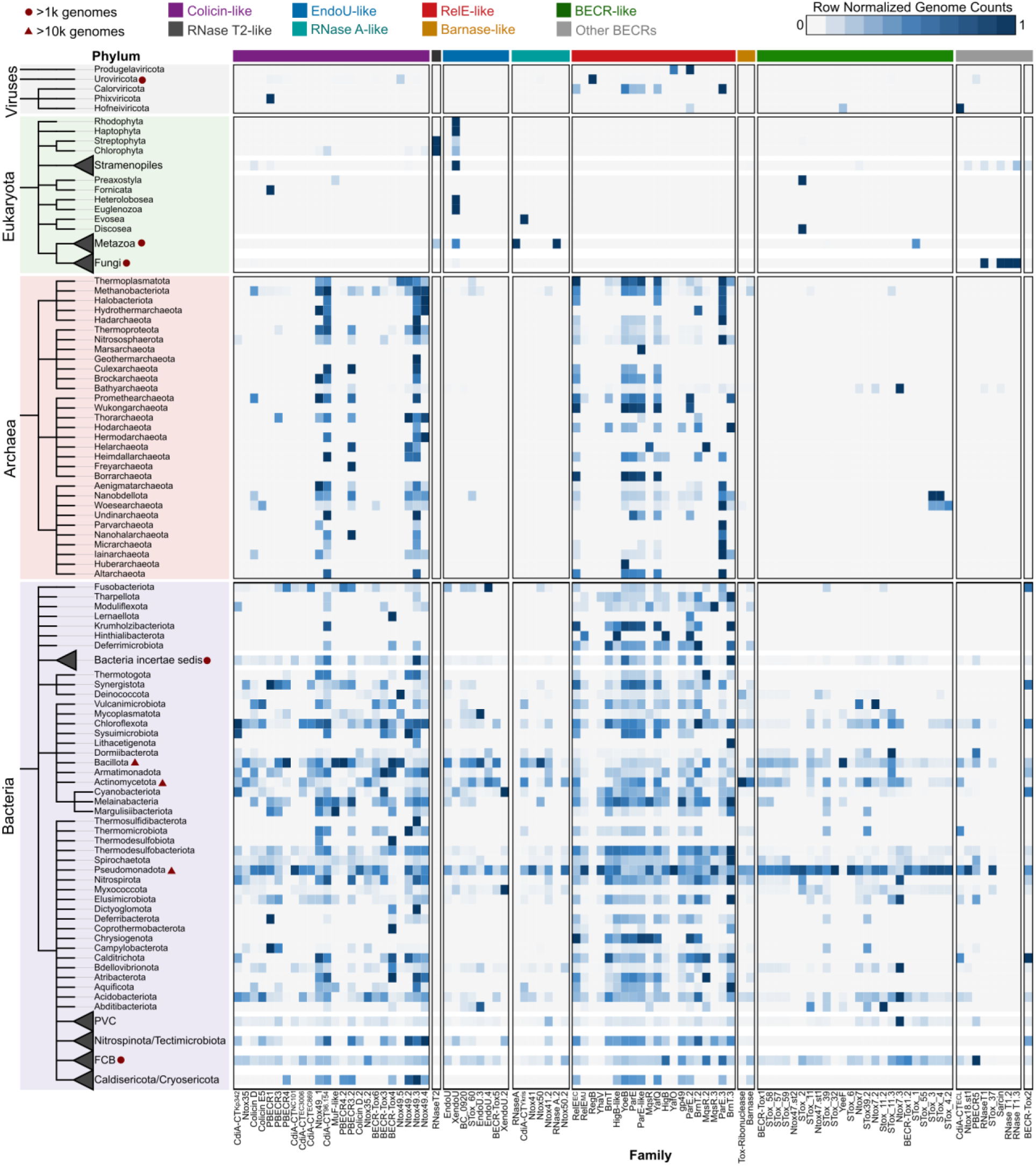
BECR RNases are widely distributed across the tree of life. A taxonomic tree collapsed at the phylum level is shown on the left, with branches colored by domain of life. Collapsed phyla are represented as isolated triangles. Along the top (x-axis), each column corresponds to a protein family, and the colored bars along the bottom indicate the assigned structural group. The heatmap displays the relative representation of each protein family across phyla, row-normalized.

The greatest diversity of BECR families was observed within the bacterial phyla Pseudomonadota, Bacillota, and Actinomycetota, which collectively harbor a large fraction of the identified toxin families. This pattern is consistent with the extensive diversification of biological conflict systems in these groups, particularly polymorphic toxin systems and other mechanisms of antagonistic interactions (Zhang et al., 2012). Gene neighborhood analyses further revealed that BECR domains occur in a wide range of conflict-associated architectures, including polymorphic toxin systems, toxin-antitoxin modules, and anti-phage defense loci (https://kristiphammie.github.io/becr/).

Notably, these associations are not uniformly distributed across the superfamily. Colicin-like proteins are frequently associated with exported antibacterial effectors, whereas RelE-like proteins are more commonly associated with intracellular toxin-antitoxin systems. Similarly, BECR-Tox2-like proteins are enriched in defense-related architectures, including Menshen-associated systems. Thus, despite sharing common structural features, different BECR lineages have been repeatedly recruited into distinct ecological and biological contexts.

Together, these results indicate that BECR RNases are both broadly distributed and evolutionarily versatile. Their uneven phylogenetic distribution, combined with their repeated deployment in diverse biological conflict systems, supports the view that the BECR fold functions as a modular platform that has been repeatedly adapted for distinct biological roles throughout evolution.

## DISCUSSION

BECR RNases illustrate how extensive sequence divergence and structural remodeling can obscure relationships among proteins that retain related biochemical functions. This problem is particularly pronounced in biological conflict systems, where toxins diversify rapidly under pressure to recognize new substrates and evade neutralization by immunity proteins. By integrating iterative homology searches, sequence- and profile-similarity networks, structural comparison, active-site mapping, and genomic context, we define an expanded organization of the BECR-fold landscape. This framework resolves major groups within the canonical BECR superfamily, identifies BECR-like proteins as a coherent but deeply divergent superfamily, and reveals previously unrecognized toxin families. Experimental validation of BECR-Tox2 further demonstrates that this guideline can identify functional toxins that remain difficult to recognize through sequence analysis alone.

The resulting organization presented here retains the canonical BECR classification established by Zhang et al. (2012) but substantially expands its membership and clarifies several previously fragmented relationships. Profile-level comparisons resolve five principal groups within the canonical superfamily: Colicin-like, EndoU-like, RelE-like, Barnase-like, and BECR-Tox2-like. They also support the proposed evolutionary connections between RNase A-like and EndoU-like proteins and between RNase T2 and Colicin-like proteins (H. Li et al., 2023; Mushegian et al., 2020). In parallel, the classification resolves ambiguous domain assignments, including the placement of DUF4258 within the Colicin-like group, the identification of DUF6883 as BECR-Tox3, and the reassignment of PF15657 to the Ntox47-related branch of the BECR-like superfamily. These examples demonstrate that family relationships obscured at the level of individual sequences can become apparent when profile similarity is interpreted together with structural topology, active-site position, and genomic context.

Across this expanded BECR landscape, the most consistently preserved feature is not the identity of individual catalytic residues or even the complete topology of the canonical scaffold. Instead, conservation is most evident in the spatial organization of the catalytic pocket relative to the structural core. Different families recruit conserved residues from distinct elements, including core β-strands, terminal extensions, duplicated modules, and lineage-specific loops. Peripheral regions, and even some elements assigned to the canonical scaffold, can be reduced without disrupting the predicted active-site geometry. In Colicin-like proteins, for example, β1 is frequently reduced or absent, whereas the active-site region formed by the pre-α element and β4 remains intact. Similarly, the duplicated N-terminal module of EndoU-like proteins varies substantially in size and organization but continues to contribute residues to a conserved catalytic arrangement. These patterns suggest that evolutionary constraints act most strongly on preservation of catalytic geometry rather than on a fixed set of residues or an invariant scaffold.

This organization provides a possible explanation for the remarkable functional diversity of BECR proteins. A structurally constrained core can maintain the geometry required for related metal-independent ribonucleolytic chemistry, whereas less constrained insertions, extensions, and surface-exposed loops can alter substrate access and interactions with protein partners. The BECR architecture therefore resembles the general principle of fold polarity, in which a relatively stable structural scaffold is coupled to more variable regions that facilitate functional diversification (Tóth-Petróczy & Tawfik, 2014). Our analyses do not directly establish that particular structural elements are more mutationally robust or evolutionarily variable but instead identify a recurrent separation between conserved catalytic organization and variable surrounding features. This model generates testable predictions, for example, family-specific elements positioned around the catalytic pocket could contribute to recognition of particular RNA substrates, while disruption of the more conserved structural core should have broader effects on catalytic activity.

These properties may also help explain why BECR domains have been repeatedly recruited into biological conflict systems. Their compact architecture, metal-independent chemistry, and capacity to tolerate extensive remodeling provide a versatile platform for generating toxins with different substrate preferences without requiring the evolution of an entirely new catalytic mechanism. Variation surrounding a conserved catalytic core could permit toxin lineages to diversify rapidly while remaining compatible with different delivery and self-protection modules. Consistent with this interpretation, BECR proteins occur in exported antibacterial toxins, intracellular toxin-antitoxin systems, antiviral defense loci, and other conflict-associated architectures. The uneven distribution of these biological roles among BECR groups further suggests that individual lineages have undergone distinct functional radiations after their incorporation into particular conflict systems. BECR-Tox2 provides a clear example of such modular deployment. Most BECR-Tox2 homologs occur in Menshen-like antiphage systems associated with a ParB-ABC ATPase module, whereas a distinct subset occurs in polymorphic toxin loci together with dedicated immunity proteins. The toxicity of BECR-Tox2^VN^ and its neutralization by BECR-Imm2^VN^ confirm the predicted toxin-immunity organization in the latter context. The occurrence of closely related BECR-Tox2 proteins in these different systems indicates that the evolutionary history of a toxin domain can be partly independent of the histories of its delivery systems and immunity proteins. A conserved catalytic domain can therefore move between biological contexts and acquire new accessory interactions without requiring an independent origin of its RNase activity.

This observation is relevant to previous interpretations of convergence among BECR toxins. Gucinski et al. (2019) described two Colicin-like CDI tRNases that possess related structures, are reciprocally detectable by sequence searches, and cleave the same tRNA isoacceptor, but differ in their immunity proteins and dependence on host factors. These differences were interpreted as evidence of convergent evolution. Our expanded analysis places both toxins within the Colicin-like group and connects them to additional families through sequence- and profile-level similarities. Their distinct accessory requirements can therefore be explained by diversification within a shared toxin lineage. More generally, the BECR-Tox2 example shows that differences in immunity proteins or genomic organization are not by themselves sufficient to distinguish independent origins from divergence after common ancestry. Convergence may still shape particular features of immunity recognition, cofactor dependence, or substrate engagement, but these properties should be interpreted separately from the evolutionary origin of the catalytic domain.

The deepest relationships within the expanded BECR landscape nevertheless remain difficult to resolve. Extreme sequence divergence precludes construction of a reliable global alignment and, consequently, a robust phylogeny capable of establishing branching order across all families. Sequence- and profile-similarity networks recover recurrent relationships among divergent groups, but network connectivity alone does not establish common ancestry. This limitation is most important for the relationship between the canonical BECR and BECR-like superfamilies. Recurrent profile-level connections, comparable organization of the predicted catalytic pocket, and deployment in similar multidomain and genomic contexts favor divergence from a common ancestral scaffold. However, the profile matches are generally weak and cover limited regions, and the small, relatively simple architecture of BECR-like proteins increases the plausibility of structural convergence. We therefore recognize BECR-like proteins as a coherent superfamily based on their shared sequence-profile, structural, and catalytic features while leaving their evolutionary relationship to canonical BECR proteins unresolved.

More broadly, the BECR landscape illustrates how substantial regions of protein space can remain fragmented when classification depends primarily on pairwise sequence similarity or existing domain models. As structure predictions become available for increasingly diverse proteins, the main challenge will shift from identifying structural resemblance to determining which similarities reflect common ancestry or convergence. The present analysis shows that no individual feature is sufficient for this task. Instead, remote relationships are most effectively evaluated through the combined evidence provided by family-level profiles, structural topology, active-site organization, and biological context.

The classification established here therefore provides both an annotation guideline and an experimental roadmap. Newly identified sequences and predicted structures can be compared against family-level combinations of profile similarity, topology, active-site position, and genomic organization rather than assigned based on a single diagnostic feature. This approach can distinguish divergent members of established families from candidate new lineages and generate testable predictions about catalytic residues, immunity partners, substrates, and biological roles. Together with the accompanying resource of structures and sequence alignments, our framework presented here transforms a fragmented collection of predicted RNases into a testable map of BECR diversity. It also establishes the BECR fold as a model for investigating how a compact catalytic core can support extensive diversification across RNA substrates and biological conflict systems.

## METHODS

### Selection of representative BECR families, structure, and sequence

Representative BECR-fold proteins were manually curated through an extensive literature review (Table S1). Families were included if they met at least one of the following criteria: (1) explicit annotation by the original authors as a BECR-fold RNase; (2) reported structural similarity to canonical BECR members; or (3) evidence of a shared ribonucleolytic cleavage mechanism. This process yielded 67 groups of sequences, referred to as “protein families” throughout this study. Initial multiple sequence alignments were retrieved for each family. When available, the multiple sequence alignment provided in the original publication or the corresponding PFAM model was used (Table S1) (Mistry et al., 2021). If no published alignment existed, we generated one by performing one iteration homology search with HHblits (Remmert et al., 2012) using default parameters against the UniRef30 database (Bateman et al., 2023). Retrieved sequences were aligned using the MAFFT local-pair algorithm (Katoh & Standley, 2013) and refined in Jalview Version 2.11.5.1 (Waterhouse et al., 2009) by removing truncated sequences. Secondary-structure predictions for each alignment were obtained using JPRED (Drozdetskiy et al., 2015). For each family, a single representative sequence and its corresponding structure were selected for analysis (Table S1 and Table S2). Representative structures were selected from experimentally determined entries in the Protein Data Bank (PDB) (Berman, 2000) whenever available. For families lacking structural data, we used high-confidence AlphaFold2 predictions (Jumper et al., 2021) already present in the database or generated new ones with ColabFold (Mirdita et al., 2022) for the representative sequence.

### Structural annotation and topology analysis

Structural annotation and visualization were performed using Mol* Viewer (Sehnal et al., 2021). For each representative structure, secondary-structure elements were compared against the corresponding family alignment to assess conservation of structural features characteristic of that family. Conservation of these elements across homologs supported their inclusion as integral components of the BECR fold. Inspection of PFAM-derived alignments revealed frequent truncations of key BECR-fold core features, most notably the N-terminal α-helix and, less commonly, the terminal β-strand. To address these inconsistencies, new sequence models were generated using the same alignment-generation workflow described above. Both the original PFAM models and the refined models produced during this analysis are provided on the website (https://kristiphammie.github.io/becr/).

### Identification of putative catalytic residues and active-site architecture

Putative catalytic residues were identified using a combination of sequence conservation and structural context. Residues were prioritized if they met the following criteria: (i) conservation across homologs within the family-specific alignment, indicating evolutionary constraint; (ii) localization within or adjacent to a putative active-site pocket, typically characterized by a partially buried or hydrophobic microenvironment; and (iii) side-chain orientation toward the interior of the pocket in structural models, consistent with a role in substrate positioning or acid–base catalysis. When available, previously characterized catalytic residues from homologous BECR families were used as additional reference points to guide interpretation (Table S1).

### Homology searches, sequence similarity network analysis, and identification of putative new families

Sequence similarity searches were initiated using the same sequences from the representative structure from each protein family as queries for transitive JackHMMER searches (Eddy, 2011) against the UniRef50 database (Suzek et al., 2007) (https://kristiphammie.github.io/becr/). JackHMMER searches were run using allocations from the Texas Advanced Computing Center (TACC) resources for five iterations using an inclusion E-value threshold of 1e-2 to maximize recovery of distant homologs. Resulting sequence sets were manually curated with the Rotifer package (https://github.com/leepbioinfo/rotifer) to remove short (<60 aa) or highly divergent sequences and to ensure retention of the core BECR-fold features and family-specific structural elements.

Redundancy reduction and clustering were performed using the MMseqs2 easy-cluster workflow (Steinegger & Söding, 2017). Searches initiated from the 67 seed sequences retrieved 69,548 homologs, which were reduced to 26,407 non-redundant sequences using clustering parameters of 0.8 coverage and 0.6 sequence identity. Sequence similarity networks (SSNs) were generated using the EFI-EST platform (Zallot et al., 2019) and visualized in Cytoscape version 3.10.4 (Shannon et al., 2003) using the yFiles organic layout (Wiese et al., 2004). Networks were constructed across a range of e-value thresholds (1e-1 to 1e-5) to evaluate the emergence of distinct clustering patterns as edge calculation parameter became more stringent. For final analyses and figures, we used the lowest alignment-score thresholds available in EFI-EST (typically 4–6) and generated representative-node networks clustered at 40% sequence identity to reduce network size and memory usage while preserving overall topological structure.

Putative new families were initially identified as discrete SSN clusters lacking a corresponding seed sequence from the curated 67 representative set (Fig. S1 and Table S2). For each of the 34 putative families, we performed additional homology searches using the same parameters applied to the initial seeds. These searches retrieved a total of 100,630 homologs, which were subsequently clustered using 0.8 coverage and a more stringent 0.5 sequence-identity threshold, yielding 22,905 non-redundant sequences. To further characterize these clusters, domain-level annotation was performed using HMMscan (Eddy, 2011) against the PFAM database to determine whether they corresponded to known protein families or represented previously unannotated groups. Additional evidence, including gene-neighborhood context, conserved motifs, and presence or absence of BECR-specific structural features, was incorporated to refine family boundaries and assess novelty.

### Profile-Profile network

Curated multiple-sequence alignments representing each protein family were compared in an all-against-all manner using HHalign from HH-suite3 (Steinegger et al., 2019). Each alignment-derived profile was represented as a node in an undirected profile–profile similarity network, and pairwise comparisons with an E-value < 1 × 10⁻³ were retained as edges. Community structure was evaluated independently using the Leiden (Traag et al., 2019) and Louvain (Blondel et al., 2008) algorithms. Network construction and analysis were performed in Python using NetworkX (Hagberg et al., 2008) and visualization in Cytoscape using Edge-weighted Spring Embedder Layout (Fig. 2).

### Gene Neighborhood Analysis

Because representative family alignments were derived from multiple databases (e.g., NCBI RefSeq/GenBank and UniProt), accession formats were first standardized to ensure consistency. For each family, HMM profiles were constructed from the curated alignments and used as queries in HMMsearch (Eddy, 2011) against the NCBI non-redundant (nr) database. Highly stringent inclusion thresholds (E-values ranging from 1e−15 to 1e−35, depending on family diversity) were applied to retrieve confident homologs while minimizing unrelated sequences. The resulting protein accessions were then used as input for either Rotifer (https://github.com/leepusp/rotifer) package or WebFlaGs (Saha et al., 2021) to retrieve and annotate gene neighborhoods.

### Toxicity Assays

*E. coli* DH5α carrying BECR-Tox2 on pBRA plasmid and BECR-Imm2 on pEXT22 plasmid were grown overnight in LB (Lysogeny Broth) media (containing 0.5% d-glucose, 50 mg/ml streptomycin, and 50 mg/ml kanamycin). Cultures were normalized to an OD₆₀₀ of 1.0 and serially diluted (1:4) in LB media. Aliquots (2 µL) of each dilution were spotted onto LB-agar plates containing streptomycin and kanamycin, supplemented with either 0.5% d-glucose or 0.2% l-arabinose and 200 µM isopropyl β-D-1-thiogalactopyranoside (IPTG). Plates were incubated at 37 °C overnight and images were acquired.

### Taxonomic Distribution

To maximize recovery of organismal diversity, all protein hits were expanded through NCBI Identical Protein Group (IPG) records rather than restricting analyses to representative genomes. Briefly, protein accessions recovered from jackHMMER searches were mapped to their associated IPG entries, and all linked genome accessions were retrieved for downstream taxonomic assignments. This strategy was chosen to avoid under sampling taxonomic diversity that can arise from representative-genome selection approaches, including those commonly used during neighborhood retrieval workflows. To reduce overrepresentation from densely sequenced species or redundant assemblies, duplicate organism entries within each family were removed prior to downstream analysis.

Taxonomic lineages were assigned using the NCBI Taxonomy database (Schoch et al., 2020) through the ete3 NCBITaxa framework. Family-level distributions were summarized across major taxonomic ranks and visualized as phylogenetic trees using ETE3 (Environment for Tree Exploration) (Huerta-Cepas et al., 2016) and the Interactive Tree of Life v6 (iTOL v6) (Letunic & Bork, 2024) platform. For comparative visualization, organism counts were aggregated across collapsed taxonomic clades and represented as associated heatmaps and abundance annotations.

## Data availability

All data supporting the findings of this study are available within the paper and its Supplementary Information files or at https://kristiphammie.github.io/becr/. The ROTIFER package can be downloaded from the GitHub repository https://github.com/leepusp/rotifer.

## Author contributions

K.P., G.G.N., R.F.S. and E.B.-S. conceived the study. K.P. performed the computational analysis with assistance from G.G.N. and R.F.S. A.I. and C.O.W. contributed to computational analyses and conceptual input. A.L performed functional assays. E.B.-S. supervised the project. All authors contributed to data analysis and interpretation. K.P, G.G.N. and E.B.-S. wrote the manuscript with input from all authors. All authors reviewed, edited, and approved the final version of the manuscript.

## Acknowledgments

This research was supported by startup funds from UT Austin College of Natural Sciences to E.B-S, Sao Paulo Research Foundation grant # 2021/10577-0 to R.F.S. C.O.W. is supported by the Blumberg Centennial Professorship in Molecular Evolution at UT Austin. The authors acknowledge the Texas Advanced Computing Center (TACC) at The University of Texas at Austin for providing computational resources that have contributed to the research results reported within this paper (http://www.tacc.utexas.edu). Additional support was provided in part by an appointment to the National Library of Medicine Research Participation Program administered by the Oak Ridge Institute for Science and Education (ORISE) though an interagency agreement between the U.S. Department of Energy (DOE) and the National Library of Medicine. ORISE is managed by ORAU under DOE contract number DE-SC0014664. All opinions expressed in this paper are the author’s and do not necessarily reflect the policies and views of NIH, NLM, DOE, or ORAU/ORISE. This work is supported by the funds of the Intramural Research Program of National Library of Medicine at the National Institutes of Health, USA (to L.A. and G.G.N). This work utilized the NIH HPC Biowulf computer cluster.

## SUPPLEMENTARY MATERIAL

**Fig. S1. Initial sequence similarity network reveals candidate new BECR families.** Sequence similarity network constructed from the first 69,548 homologs, used to identify putative new BECR families. Clusters lacking any of the 67 initial seed sequences were flagged as candidate new families. Edges were calculated using an e-value threshold of 1e-2. Black nodes represent the original seed sequences, and red nodes denote newly identified seed sequences.

**Fig. S2. AlphaFold-Multimer analysis supports interaction between BECR-Tox2 and BECR-Imm2.** (A) Predicted Aligned Error (PAE) and predicted Local Distance Difference Test (pLDDT) confidence scores from the AlphaFold-Multimer prediction. (B) Predicted complex of BECR-Tox/Imm2. BECR-Tox2 (grey) with conserved residues shown in Fig 5A as opaque molecular surface representation, and BECR-Imm2 (orange) with residues marked in (C) shown as yellow ball-and-stick. Right panel shows zoomed in view of interacting Y20 from BECR-Imm2. (C) Sequence logo for BECR-Imm2 highlighting conserved residues denoted by yellow markers.

**Table S1. Comprehensive list of all BECR proteins used in this study.**

**Table S2. Comprehensive sequence datasets used throughout this study.** (A) Initial 13,434 BECR sequences compiled from publications, PFAM entries, and original alignments generated in this study. (B) Curated representative seed set for the 67 initial BECR families selected from the original alignments. (C) 69,547 homologs retrieved from first JackHMMER searches for the 67 initial seeds. (D) Representative clustered dataset for SSN construction including 29,407 sequences obtained after clustering at 0.8 coverage and 0.6 identity, corresponding to the network shown in Figure S1. (E) Seed sequences for 34 putative new BECR families selected from clusters lacking initial seeds, as highlighted in red in Figure S1. (F) Combined homolog dataset for expanded family searchers - 100,630 total homologs including sequences from initial and putative new families. (G) Final representative clustered dataset for global analyses containing 22,905 sequences obtained after clustering at 0.8 coverage and 0.5 identity, corresponding to Fig 3B.

