## Supplementary figures and images for "A structure-guided classification framework reveals the diversity and catalytic architecture of BECR ribonuclease"

### Fig. S1

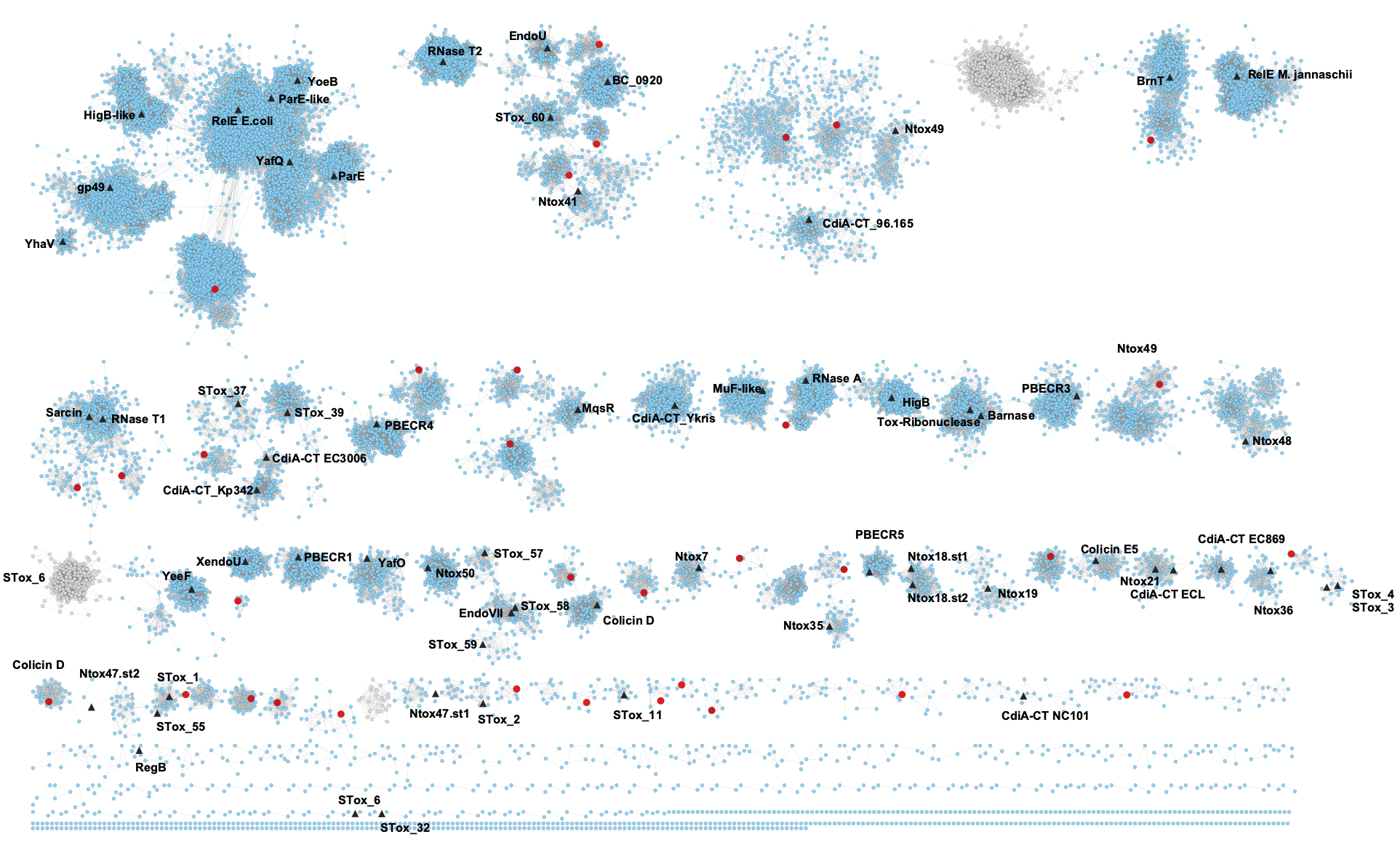

### Fig. S2

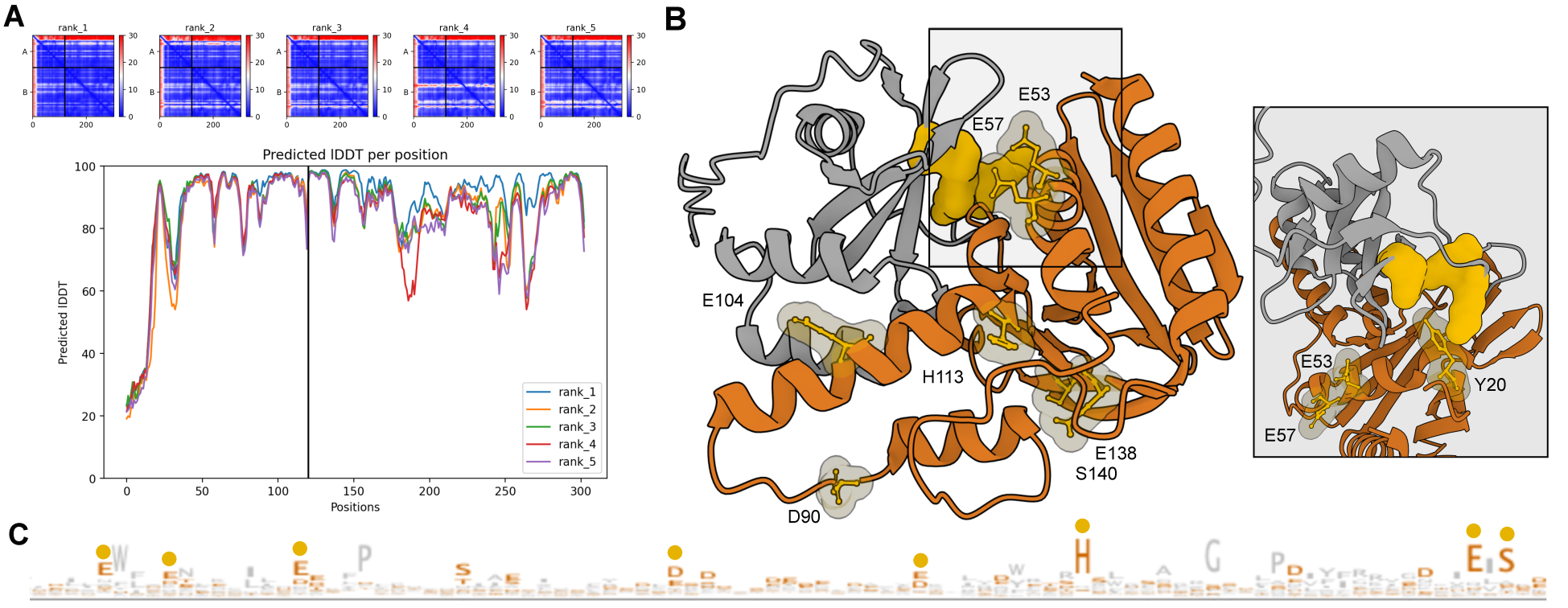
